# A Jacalin-like lectin-domain-containing protein of *Sclerospora graminicola* act as an apoplastic virulence effector in plant–oomycete interactions

**DOI:** 10.1101/2021.08.30.458171

**Authors:** Michie Kobayashi, Hiroe Utsushi, Koki Fujisaki, Takumi Takeda, Tetsuro Yamashita, Ryohei Terauchi

## Abstract

The plant extracellular space, including the apoplast and plasma membrane, is the initial site of plant– pathogen interactions. Pathogens deliver numerous secreted proteins, called effectors, into this region to suppress plant immunity and establish infection. Downy mildew caused by the oomycete pathogen *Sclerospora graminicola* (Sg) is an economically important disease of Poaceae crops including foxtail millet (*Setaria italica*). We previously reported the genome sequence of Sg and showed that the Jacalin-related lectin (JRL) gene family has significantly expanded in this lineage. However, the biological functions of JRL proteins remained unknown. Here, we show that JRL from *S. graminicola* (SgJRL) functions as an apoplastic virulence effector. We identified eight SgJRLs *via* protein mass spectrometry analysis of extracellular fluid from *S. graminicola*-inoculated foxtail millet leaves. SgJRLs consist of a Jacalin-like lectin domain and an N-terminal putative secretion signal, and *SgJRL* expression is induced by Sg infection. Heterologous expression of three SgJRLs with N-terminal secretion signal peptides in *Nicotiana benthamiana* enhanced the virulence of the pathogen *Phytophthora palmivora* inoculated onto the same leaves. Of the three SgJRLs, SG06536 fused with GFP localized to the apoplastic space in *N. benthamiana* leaves. INF1-mediated induction of defense-related genes was suppressed by co-expression of SG06536-GFP. These findings suggest that JRLs are novel apoplastic effectors that contribute to pathogenicity by suppressing plant defense responses.

## INTRODUCTION

To counter the constant attacks by pathogens, plants mount defense responses (Jones and Dangl, 2006). The first battleground of plants and pathogens is the extracellular space, including plant surfaces and the apoplast. Plants release a battery of hydrolases, protease inhibitors, and antimicrobial compounds into the apoplast to prevent microbial infection (Doehlemann and Hemetsberger, 2013; Wang and Wang, 2018; Wang et al, 2019). Plant proteases and protease inhibitors suppress pathogen growth directly or indirectly (Jashni et al, 2015). In addition, plant β-1,3-glucanase and chitinases degrade major components of the cell walls of fungi and oomycetes. Pattern recognition receptors (PRRs) localized on the plant cell surface sense elicitor molecules such as pathogen-associated molecular patterns (PAMPs), microbe-associated molecular patterns (MAMPs), and damage-associated molecular patterns (DAMPs) and activate immune responses. As a counter-defense, pathogens manipulate plant immune responses using virulence effectors, resulting in successful infection.

Oomycetes (also known as water molds) are a diverse group of filamentous eukaryotic microorganisms that include saprophytes as well as pathogens of plants, insects, crustaceans, fish, vertebrate animals, and various microorganisms (Lamour and Kamoun 2009, Thines and Kamoun, 2010, Wang et al, 2019). More than 60% of known oomycete species are parasitic on plants (Thines and Kamoun, 2010), in which they cause devastating diseases in a wide range of species, including agricultural crops. Oomycetes secrete a series of effectors to manipulate plant physiology and suppress plant immunity (Kamoun, 2006). Whole-genome sequencing and transcriptome analysis of various oomycetes revealed that they share a common set of effectors. These effectors are classified as apoplastic or cytoplasmic based on their localization in the host plant. Apoplastic effectors include secreted hydrolytic enzymes such as proteases, lipases, and glycosylases that can degrade plant tissue, protease inhibitors that protect oomycetes from host defense enzymes, necrosis and ethylene-inducing peptide 1 (Nep1)-like proteins (NLPs), and PcF-like small cysteine-rich proteins (SCRs) (Kamoun, 2006). By contrast, RXLR-domain-containing proteins and crinklers (CRNs) are cytoplasmic effectors that are specific to plant pathogenic oomycetes (Torto et al, 2003, Morgan and Kamoun 2007).

*Sclerospora graminicola* (Sacc.) Schroet. is an obligate biotrophic oomycete that causes downy mildew disease in Poaceae plants including foxtail millet, pearl millet, and maize. We previously performed whole-genome sequencing of *S. graminicola* and determined that a gene family encoding Jacalin-like lectin-domain containing proteins (JRLs) significantly has expanded in its genome (Kobayashi et al, 2017). Many SgJRLs have putative secretion signals at their N-termini and their expression is upregulated during *S. graminicola* infection, suggesting that SgJRLs are novel effectors of *S. graminicola*. However, the functions of SgJRLs remain unknown, primarily due to the difficulty of genetically manipulating oomycetes, including *S. graminicola*.

Here we report that JRLs of *S. graminicola* are indeed secreted into the apoplast. Using the model pathosystem of *Nicotiana benthamiana* and *Phytophthora palmivora*, we show that transient overexpression of *SgJRL* in the plant enhances *P. palmivora* growth and partially suppresses plant defense responses. These findings indicate that JRL is a novel apoplastic effector that contributes to the virulence of oomycetes.

## RESULTS

### *S. graminicola* delivers JRLs into the plant apoplast during early infection

To investigate the molecular mechanisms underlying plant–microbe interactions in the apoplastic space during *S. graminicola* infection, we extracted extracellular fluid (EF) from *S. graminicola*-inoculated foxtail millet leaves at 1 and 3 days post-inoculation (dpi). We then compared the proteins contained in the EF to those of the soluble fraction (SF) from total leaf homogenates. Thaumatin-like protein, an apoplastic marker protein, was detected by anti-thaumatin antibody in the EF but not in the SF, whereas FBPase, a cytoplasmic marker as detected by anti-FBPase antibody, was abundant in the SF but not in the EF (Fig. S1), indicating that apoplastic proteins were considerably enriched in our EF preparation. Therefore, we subjected the EF proteins to mass spectrometry analysis (Fig. S2). LC-MS/MS analysis identified 75 *S. graminicola* proteins, including 19 putative secreted proteins (Table S1). Most putative secreted proteins were contained in the low-molecular-weight fraction from leaves at 3 dpi (Table S1, fractions 3dpi-3 to 3dpi-5). Therefore, we performed another experiment focusing on the low-molecular-weight fractions, which identified 23 *S. graminicola* proteins, including 10 putative secreted proteins (Table S2). Seven putative secreted proteins were common to the first and second experiments (Table 1, shaded).

**Table 1.** Putative secreted proteins of *Sclerospora graminicola* identified in the extracellular fluid of *Sclerospora graminicola*-inoculated foxtail millet leaves.

| Accession | Mass | Descriptions | Exp_1<sup>a</sup> | Exp_2<sup>a</sup> |
| --- | --- | --- | --- | --- |
| SG14735 | 35,549 | arabinofuranosidase | ✓ |  |
| SG08752 | 49,348 | calreticulin precursor | ✓ |  |
| SG02395 | 50,987 | exonuclease | ✓ |  |
| SG14267 | 61,637 | glucose-methanol-choline oxidoreductase | ✓ |  |
| SG02945<sup>b</sup> | 19,042 | JRL | ✓ | ✓ |
| SG03255<sup>b</sup> | 16,328 | JRL | ✓ | ✓ |
| SG04896 | 18,357 | JRL |  | ✓ |
| SG05109<sup>b</sup> | 19,699 | JRL | ✓ | ✓ |
| SG05127 | 18,563 | JRL | ✓ |  |
| SG06536<sup>b</sup> | 16,412 | JRL | ✓ | ✓ |
| SG10124<sup>b</sup> | 17,422 | JRL | ✓ | ✓ |
| SG17166<sup>b</sup> | 18,094 | JRL | ✓ | ✓ |
| SG00344 | 26,908 | NLP | ✓ |  |
| SG05755<sup>b</sup> | 29,163 | NLP | ✓ | ✓ |
| SG10403 | 27,419 | NLP | ✓ |  |
| SG14292 | 26,664 | NLP |  | ✓ |
| SG05676 | 9,704 | phospholipase D | ✓ |  |
| SG14096 | 13,994 | protease inhibitor | ✓ |  |
| SG18945 | 640,765 | protocadherin fat-like protein | ✓ |  |
| SG12053 | 45,595 | transglutaminase elicitor | ✓ |  |
| SG00883 | 22,153 | uncharacterized | ✓ |  |
| SG03074 | 18,464 | uncharacterized |  | ✓ |

a: Exp\_1 and Exp\_2 indicate proteins identified in first and second experiments, respectively.
b: Proteins identified in both experiments.

Combining the two experiments, a total of 22 proteins were identified as putative apoplastic secreted proteins, including eight Jacalin-related lectin proteins (JRLs), four necrosis and ethylene-inducing peptide 1 (Nep1)-like proteins (NLPs), one arabinofuranosidase, one protease inhibitor, and one transglutaminase elicitor, among others (Table 1). Phylogenetic analysis and amino acid sequence alignment of the SgJRLs showed that the eight JRLs are not similar to each other among SgJRL family members, but all contained a Jacalin domain with conserved amino acid residues (Fig. 1 and Fig. S3). All eight *JRL* genes were induced during *S. graminicola* infection, with their highest expression at 2 dpi (Fig. 2). These data indicate that JRLs are candidate apoplastic effectors with possible roles in virulence during the early infection stage (∼2 dpi). Therefore, we focused on the JRLs in subsequent analysis.

**Fig. 1.**
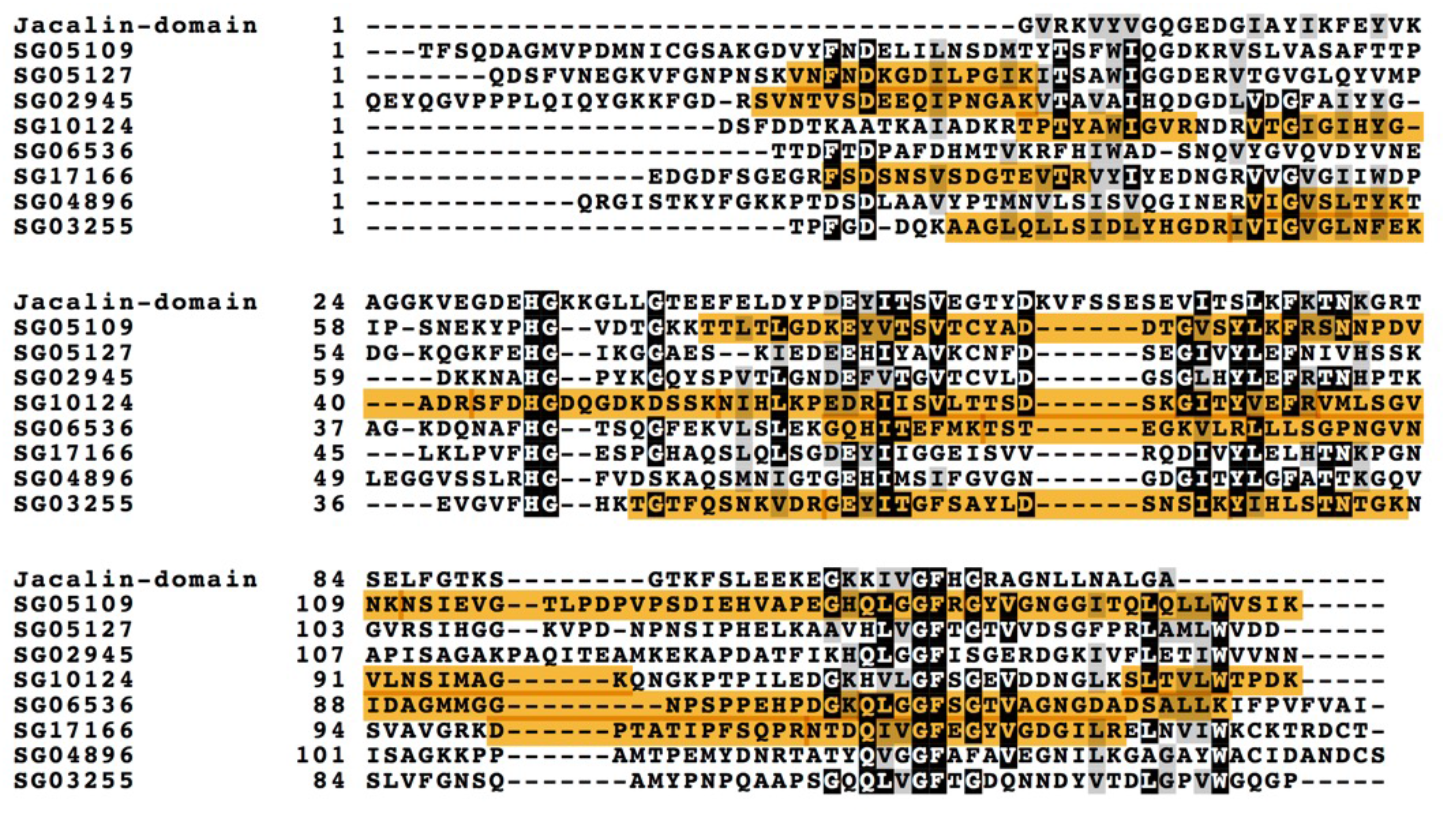
Amino acid sequence alignment of the Jacalin-like lectin domain consensus sequence (Jacalin-domain) and SgJRLs without the N-terminal signal sequence. Multiple sequence alignment was performed using ClustalW and illustrated by BOXSHADE (https://embnet.vital-it.ch/software/BOX_form.html). Identical amino acids are shaded in black, and similar amino acids are shaded in gray. Orange shading indicates peptides identified by LS-MS analysis.

**Fig. 2.**
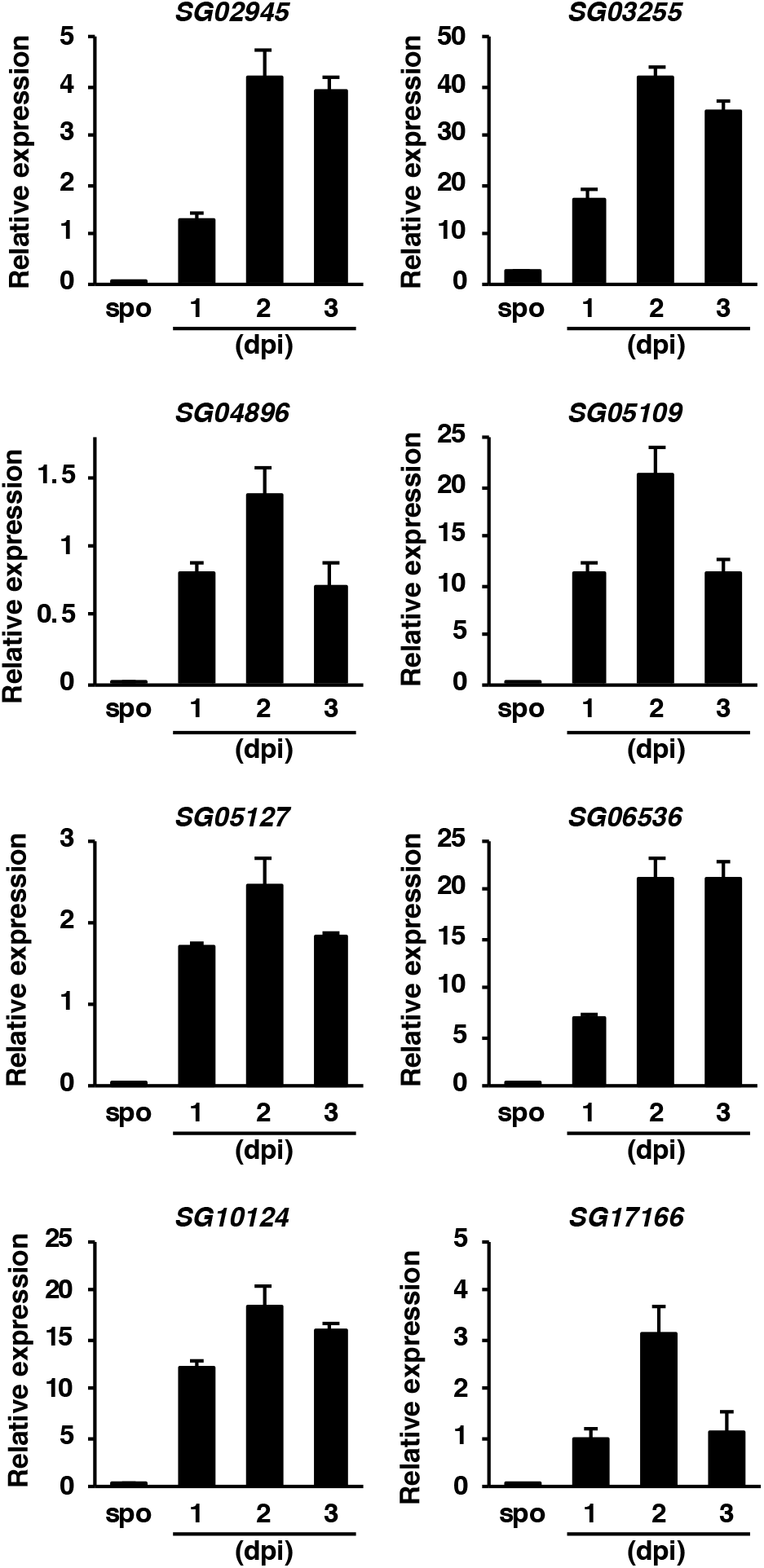
Relative expression levels of genes encoding JRLs identified in the extracellular fluid from foxtail millet leaves inoculated with *S. graminicola. Histone H2A* was used as a reference gene for normalization. spo, *S. graminicola* conidiospores. (*n* = 3.)

### JRLs with secretion signals promote *P. palmivora* infection in *N. benthamiana* leaves

Since genetic manipulation of foxtail millet and *S. graminicola* is difficult, we used a model system consisting of *N. benthamiana* and *P. palmivora* to evaluate the possible virulence functions of JRLs. To investigate the functions of SgJRLs in the apoplast, we replaced their predicted secretion signal peptides with that of *Nicotiana tabacum* PR1a (NtPR1a; GenBank accession: X06930.1) (Fig. 3A). Since the NtPR1a signal peptide is highly effective at targeting proteins to the apoplast *via* the endoplasmic reticulum (ER)/Golgi secretory pathway in solanaceous plants (Hammond-Kosack et al., 1994), we expected that SgJRLs with the NtPR1a signal peptide would accumulate in the apoplast. To test the effects of JRL expression, we transiently expressed eight JRLs (designated SG02945, SG03255, SG04896, SG05109, SG05127, SG06536, SG10124, and SG17166 in this study) with or without the NtPR1a signal peptide in *N. benthamiana* leaves and inoculated the same leaves with *P. palmivora* 2 days after agroinfiltration. At 3 dpi, transient expression of the three proteins (SG03255, SG06536, and SG17166) with signal peptides resulted in larger disease lesion sizes than transient expression of their counterpart proteins lacking signal peptides (Fig. 3B). Expressed SG03255, SG06536, and SG17166 proteins of the predicted sizes were detected by immunoblot analysis (Fig. 3C). These results suggest that JRL proteins SG03255, SG06536, and SG17166 are secreted into the apoplast and have virulence functions that enhance *P. palmivora* growth in planta.

**Fig. 3.**
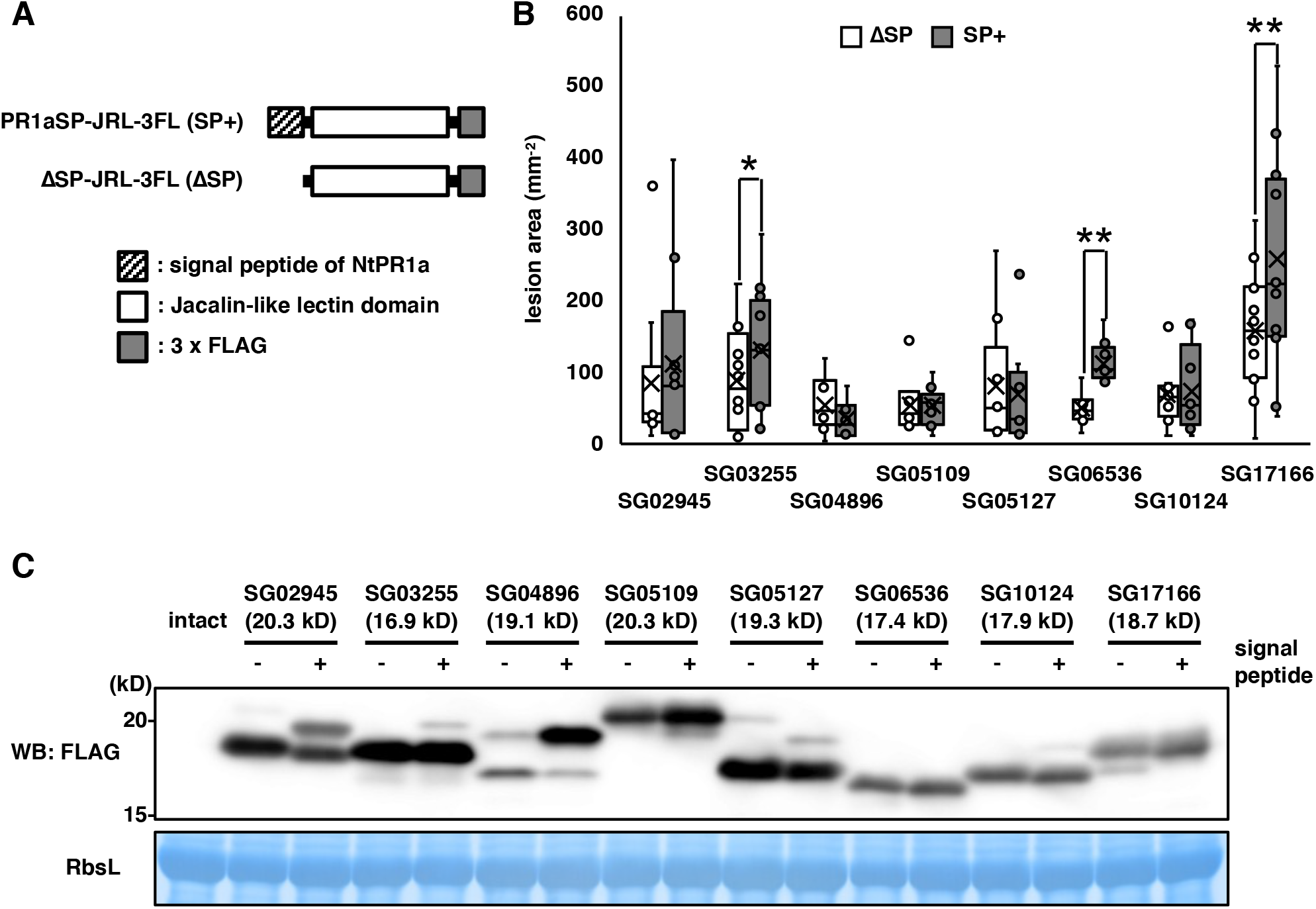
JRLs with secretion signals promote *P. palmivora* infection in *N. benthamiana* leaves. (A) Diagrams of SgJRL and the fusion proteins used in this study. (B) JRLs with or without the secretion signal sequence from NtPR1a were transiently expressed in *N. benthamiana* leaves. Two days after agroinfiltration, the leaves were detached and inoculated with *P. palmivora*. Lesion area was measured at 3 dpi. Asterisks indicate significant differences (*, *P* < 0.05; **, *P* < 0.01; *n* = 8–12) using paired two-tailed Student’s *t* test. (C) Leaves expressing JRLs were sampled 2 days after agroinfiltration, and proteins were detected using anti-FLAG antibody.

### SP-SG06536-GFP localizes to the apoplast in *N. benthamiana* leaves

To validate the localization of the SgJRLs to the apoplast, we fused GFP to the C-terminus of each SgJRL with or without the secretion signal from NtPR1a (PR1aSP-SgJRL-GFP and ΔSP-SgJRL-GFP, respectively) (Fig. 4A, Fig. S4A). We then transiently expressed the PR1aSP-SgJRL-GFP and ΔSP-SgJRL-GFP fusion proteins in *N. benthamiana* leaves and visualized them by confocal microscopy. By combined plasmolysis and FM4-64 staining, we detected the accumulation of PR1aSP-SgJRL-GFP in the apoplastic space, while the ΔSP-SgJRL-GFP proteins and the control (GFP) remained in the cytoplasm, except for SG17166 (Fig. 4B, Fig. S4B). These observations confirm the translocation of PR1aSP-SgJRL-GFP to the region outside the cells and imply that it functions in the apoplast.

**Fig. 4.**
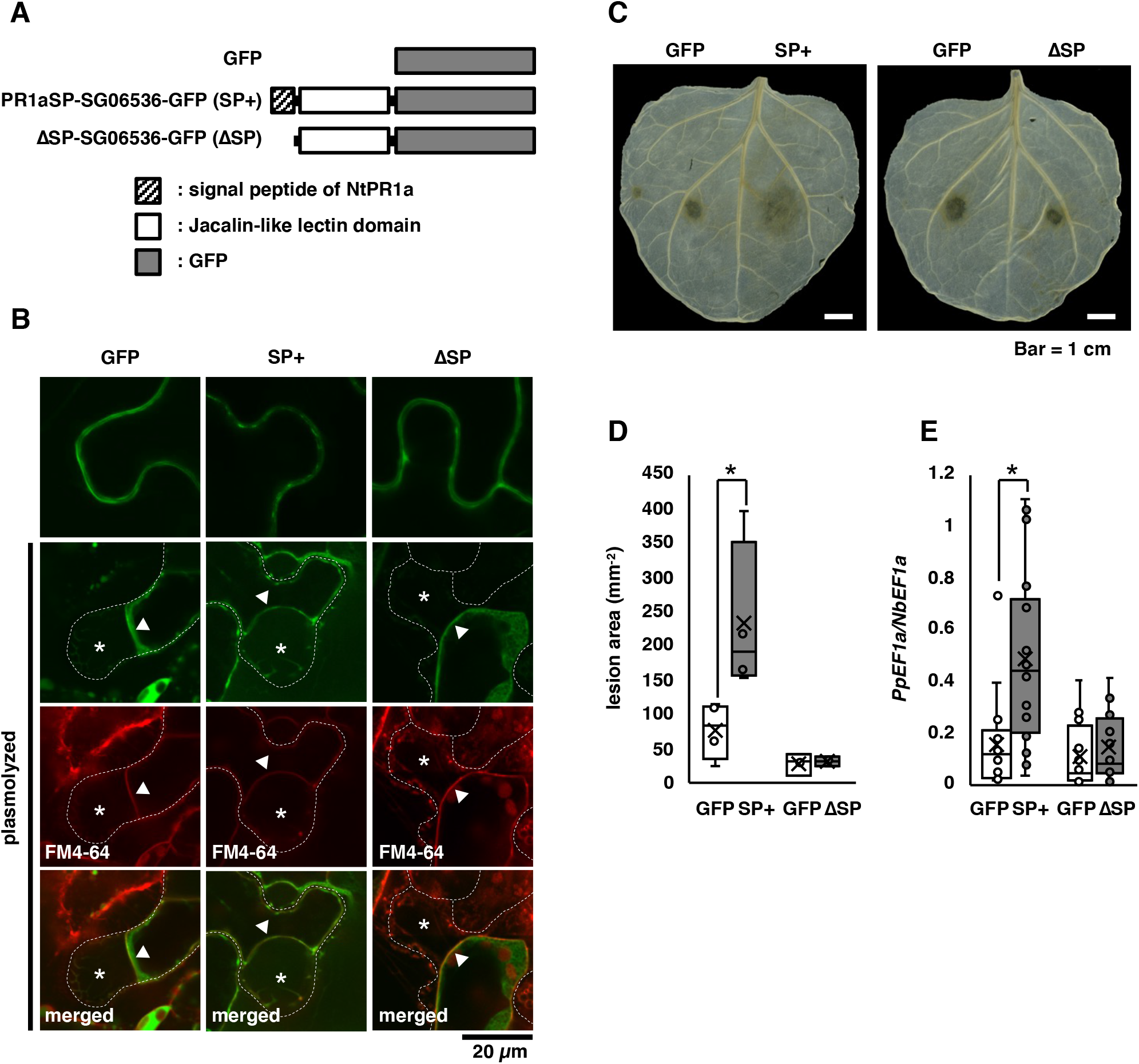
SG06536 with a signal peptide accumulates in the plant apoplast. (A) Diagrams of SG06536 and the fusion proteins used in this study. (B) Genes were transiently expressed in *N. benthamiana* leaves. The leaves were examined 3 days after agroinfiltration. In plasmolyzed cells, the dashed white line marks the cell wall, asterisks indicate the apoplastic space, and triangles indicate the position of the plasma membrane. Bar = 20 µm. (C) SG06536 with or without the secretion signal sequence from NtPR1a was transiently expressed in *N. benthamiana* leaves. Two days after agroinfiltration, the leaves were detached and inoculated with *P. palmivora*. The leaves were decolored and photographed at 3 dpi. (D) Lesion area was measured at 3 dpi. Asterisk indicates a significant difference (*P* < 0.05, *n* = 3 or 4) using paired two-tailed Student’s *t* test. (E) The ratio of *P. palmivora*/*N. benthamiana* biomass assayed by qPCR using DNA isolated from leaves at 3 dpi. Asterisk indicates a significant difference (*P* < 0.05, *n* = 16–18) using paired two-tailed Student’s *t* test.

Of the JRLs examined, the effect of SG06536 on *P. palmivora* growth showed the clearest difference in the presence versus the absence of the signal peptide (Fig. 3B). Therefore, we evaluated the growth of *P. palmivora* in *N. benthamiana* overexpressing SG06536-GFP. Compared to GFP, PR1aSP-SG06536-GFP but not ΔSP-SG06536-GFP promoted the spread of *P. palmivora* disease lesions in *N. benthamiana* (Fig. 4C, D). In addition, the biomass of *P. palmivora* was correlated with the sizes of the disease lesions (Fig. 4E). Transiently expressed PR1aSP-SG06536-GFP and ΔSP-SG06536-GFP proteins accumulated to similar levels in *N. benthamiana* leaves, as determined by immunoblot analysis (Fig. S5). These data are consistent with the results above (Fig. 3B) and highlight the importance of the secretion of SG06536 into the apoplast to support *P. palmivora* infection.

### SG06536 expression inhibits PAMP-triggered responses

To investigate the effects of SG06536 on plant defense responses, we examined the expression of defense-related genes *PAL* and *RBOHB*. INF1 is a PAMP derived from *Phytophthora* species (Kamoun et al. 1997) that induces plant defense responses. In agroinfiltrated leaves, *PAL* expression was induced and reached a peak at 9 h after treatment, and *RBOHB* expression increased until 12 h after treatment (Fig. S6). We then infiltrated INF1 peptides into *N. benthamiana* leaves expressing *GFP, PR1aSP-SG06536-GFP*, and *ΔSP-SG06536-GFP* at 2 days after agroinfiltration. At 9 h after INF1 treatment, the expression levels of *PAL* and *RBOHB* were lower in *PR1aSP-SG06536-GFP-*expressing leaves than in *GFP*- and *ΔSP-SG06536-GFP*-expressing leaves (Fig. 5). These results suggest that SG06536 in the apoplastic space suppresses PAMP-triggered responses.

**Fig. 5.**
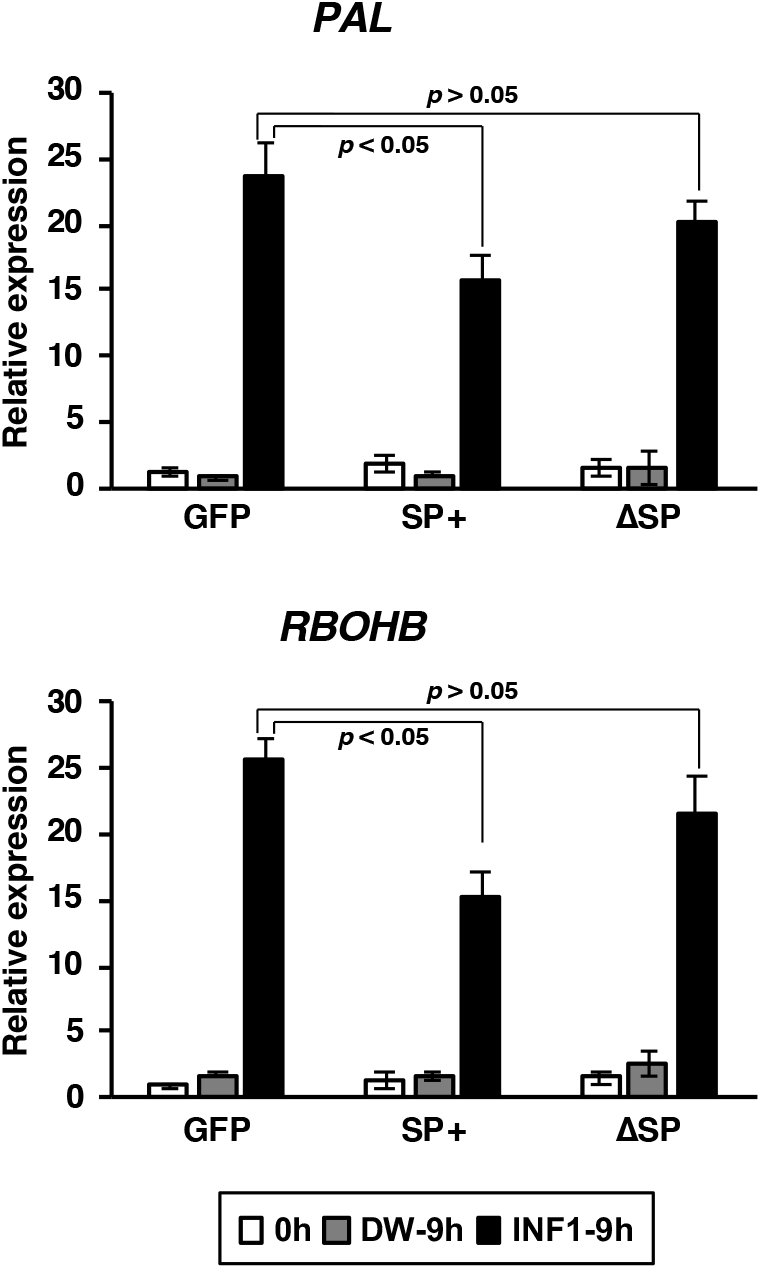
INF1-mediated induction of *PAL* and *RBOHB* in *N. benthamiana* leaves is partially suppressed by the expression of SP-SG06536-GFP. Two days after agroinfiltration, the leaves were subjected to INF1 treatment. The relative expression levels were analyzed by qRT-PCR, with *NbEF1α* used as a reference gene for normalization. *P*-values were calculated using Dunnett’s multiple-comparisons test.

## DISCUSSION

In this study, we investigated *S. graminicola* proteins in the host apoplast. Although the purity of the apoplastic fraction is often a concern in the analysis of apoplastic proteins, the EF produced in this study appeared to be significantly enriched with apoplastic proteins, as demonstrated by immunoblot analysis of thaumatin-like protein, an apoplastic marker (Fig. S1). Supporting this notion, LC-MS/MS analysis identified several plant apoplastic proteins in the EF, including PR proteins with high scores than that in the SF (Tables S4 and S5). We also identified several *S. graminicola* apoplastic effector candidates in the EF (Tables S1 and S2). NLPs are widespread effectors among filamentous and bacterial pathogens. In oomycetes, two types of NLPs (type 1 : cytotoxic-type and type 1a : non-cytotoxic-type) have been reported (Oome et al, 2014). The SgNLPs identified in this study are type 1a NLPs; such NLPs are thought to play a role in the biotrophic phase of oomycete infection (Oome et al, 2014), but the details of their function remain unclear. Further analysis of SgNLPs will be required to understand the infection strategy of *S. graminicola*.

In the current study, we focused on SgJRLs, since the *SgJRL* gene family appears to have undergone a unique expansion process in the *S. graminicola* genome among related oomycetes (Kobayashi et al, 2017). The *S. graminicola* genome encodes 45 JRLs with putative N-terminal secretion signals. Eight of these proteins were identified in our LC-MS/MS analysis in the current study. All eight SgJRLs were highly expressed during the early phase of *S. graminicola* infection, which is in agreement with their abundance detected by MS analysis (Fig. 2, Tables S1 and S3). Three of these eight (SG03255, SG06536, and SG17166) promoted *P. palmivora* infection in *N. benthamiana* leaves (Fig. 3B). Phylogenetic analysis and amino acid sequence alignment of the SgJRLs showed that the three JRLs are not similar to each other among SgJRL family members (Fig. 1 and Fig. S3). Therefore, we cannot infer the functions of JRLs based on their primary sequences. Determining the three-dimensional structures of the SgJRLs would be useful for elucidating their functions in the future (de Guillen et al, 2015, Win et al, 2012).

JRLs were previously identified as candidate effectors based on genome analysis of *Phytophthora infestans* (Raffaele et al, 2010). However, the functions of JRLs in filamentous pathogens are not known. JRLs are lectins that reversibly bind to carbohydrates such as mannose and galactose. Several pathogen carbohydrate-binding proteins function as virulence effectors in the apoplast. For example, the LysM-domain-containing proteins Ecp6, Slp1, and Mg3LysM prevent chitin-triggered plant immunity by binding to chitin oligosaccharides (de Jonge et al, 2010, Mentlak et al, 2012, Marshall et al, 2011). Mg1LysM, Mg3LysM, and Avr4 protect fungal hyphae against plant-derived hydrolytic enzymes, probably by binding to the fungal cell wall (Marshall et al, 2011, van den Burg et al, 2006). The fungi-specific β-glucan-binding lectin FGB1 alters fungal cell wall composition to protect against plant hydrolases, and it also suppresses β-glucan-triggered immunity (Wawra et al, 2016). All of these factors contribute to fungal pathogenicity by binding to carbohydrates derived from fungal cell walls. We have not yet successfully detected the binding of SgJRLs to the cell walls of either oomycetes or foxtail millet. NLP, which has structural similarities to fungal lectins, binds to glycosylinositol phosphorylceramide (GIPC), a major lipid of the plant plasma membrane, and causes necrosis in eudicots (Lenarčič et al, 2017; Ottmann et al, 2009). These observations point to possible roles of JRLs in the apoplast.

In the current study, we identified three JRLs (SG03255, SG06536, and SG17166) that affect *P. palmivora* infection in *N. benthamiana*, two of which (PR1aSP-SG03255-GFP and PR1aSP-SG06536-GFP) accumulate in the apoplastic space, although PR1aSP-SG17166-GFP was not visibly localized to the apoplast (Fig. S4). We also demonstrated that PR1aSP-SG06536-GFP expression has an inhibitory effect on plant defense responses (Fig. 5). These data suggest that at least the secreted protein SG06536 supports *P. palmivora* infection and suppress plant defense responses in the apoplast.

In contrast to JRLs from pathogens, information is available about the roles of plant JRLs in disease resistance (Esch and Schaffrath 2017). *TaJRLL1*, a wheat JRL, is induced by biotic and abiotic stimuli and is thought to be a component of salicylic-acid- and jasmonic-acid-dependent defense signaling (Xiang et al, 2011). Overexpression of *OsJAC1* confers quantitative broad-spectrum resistance against fungal, oomycete, and bacterial pathogens in rice (*Oryza sativa*; Weidenbach et al, 2016). Although they are likely to be intracellular proteins, some host JRLs can function in the apoplast: for example, rice OsMBL1. Overexpression of *OsMBL1* induces resistance to rice blast (caused by *Magnaporthe oryzae*), whereas MoChi1, an *M. oryzae* chitinase, inhibits the resistance response by competing with OsMBL1 for chitin (Han et al, 2019). Therefore, it is possible that SgJRLs mimic or compete with host JRLs that function in the apoplast.

We demonstrated that an SgJRL functions as a novel virulence effector. Genome analysis showed that oomycetes closely related to *S. graminicola* have JRLs, whereas *Pythium* spp., an oomycete distantly related to *S. graminicola* and other fungi, such as *M. oryzae* and *Colletotrichum graminicola*, do not have low-molecular-weight JRLs with signal peptides encoded in their genomes (Table S6). Notably, *S. graminicola* has an uniquely expanded JRL family among related oomycetes (Kobayashi et al, 2017). Among the oomycetes with available genome sequences, only *S. graminicola* infects monocots. Therefore, perhaps the expansion of the JRL family represents an adaptation of this pathogen to monocotyledons. The differential functions of JRLs in monocotyledons and dicotyledons are of great interests and should be clarified in the future.

## EXPERIMENTAL PROCEDURES

### Plant and oomycete materials and inoculation

Foxtail millet (*Setaria italica* (L.) P. Beauv., cultivar ‘Ootsuchi-10’), obtained from the experimental field of Iwate Agricultural Research Center (IARC), Karumai, Iwate, Japan, with permission, was used in this study. A strain of *Sclerospora graminicola* (Sacc.) Schroet. isolated from a single zoospore was derived from an isolate collected in the IARC field with permission in 2013. The plants were grown in an artificial climate chamber at 20–25 °C under a 15 h light/9 h dark cycle. Four-week-old plants were infected with *S. graminicola* by spraying them with a mixture of sporangia and zoospores (1–5 × 10^5^ per mL). Seven days after inoculation, the leaves were harvested, rinsed with distilled water, and used to prepare the inoculum. Sporulation was induced by incubating the infected leaves at 100% humidity at 25 °C for 5–6 h. Mixtures of sporangia and zoospores were collected by rinsing the sporulated leaves with chilled sterile water. *N. benthamiana* plants were grown for 4–5 weeks at 25 °C under a 16 h light/8 h dark cycle. *P. palmivora* (MAFF 242760) was maintained on rye medium at 25 °C. For inoculation, *P. palmivora* was subcultured on V8 medium in the dark for 3 days, followed by 3 days under a 16 h light/8 h dark cycle. The mycelium was suspended in cold water, filtered through Miracloth and concentrated by centrifugation, and zoospore production was induced at room temperature. Detached *N. benthamiana* leaves were inoculated with 10-µl aliquots of zoospores (1 × 10^5^ zoospores/ml) and covered with lens paper (4 mm in diameter) to keep the zoospore suspension on the leaf surface. The leaves were kept at high humidity at 25 °C.

### Preparation of the extracellular fluid and soluble protein fractions

Foxtail millet leaves inoculated with *S. graminicola* were harvested and vacuum-infiltrated with 50 mM HEPES-KOH, pH 6.8, for 20 min. The leaf surface was dried, and the leaf material was placed into a 25-mL syringe and inserted into a 50-mL Falcon tube, which was then centrifuged at 1500 × *g* for 10 min. An aliquot of the sample was immediately mixed with protease inhibitor cocktail. The sample was used as the extracellular fluid (EF) fraction for mass spectrometry and immunoblot analysis. To extract the soluble protein fraction, the leaves were ground in liquid nitrogen and dissolved in three volumes of GTN extraction buffer (25 mM Tris-HCl, pH 7.0, 150 mM NaCl, 10% glycerol, 10 mM dithiothreitol, and complete protease inhibitor cocktail). The samples were centrifuged at 20,000 × *g* for 10 min and the supernatant used as the soluble protein fraction (SF).

### Mass spectrometry

The proteins were analyzed by SDS-PAGE and CBB staining (Fig. S1 and S2). The protein bands were excised from the gel, treated with trypsin, and analyzed by mass spectrometry using an LTQ Orbitrap XL mass spectrometer (Thermo Fisher Scientific, Inc., USA) as described previously (Takahashi et al., 2013).

### *Agrobacterium tumefaciens*-mediated transient expression (agroinfiltration) in *N. benthamiana*

Binary plasmids were transformed into Agrobacterium strain GV3101 and cultured overnight. The culture was diluted 10-fold in Luria-Bertani medium containing kanamycin and rifampicin and cultured until it reached OD_600_ = 0.6. The cells were harvested by centrifugation and resuspended in MES/MgCl_2_ (10 mM MES-NaOH, pH 5.6, 10 mM MgCl_2_, 150 µM acetosyringone). The suspensions were adjusted to OD_600_ = 1.0 and incubated for 1–2 h at room temperature. Bacterial suspensions containing P19, SgJRL, and MES/MgCl_2_ were mixed at a 1:5:4 ratio and infiltrated into the leaves of 4- to 5-week-old *N. benthamiana* plants using a needleless syringe. Two or three days after infiltration, the leaves were used for various experiments.

### Immunoblot analysis

*N. benthamiana* leaves were ground in liquid nitrogen and dissolved in six volumes of GTN extraction buffer. SDS-PAGE sample buffer was added to the samples, which were then boiled for 5 min, chilled on ice, and centrifuged at 20,000 × *g* for 10 min to remove debris. Equal volumes of samples were separated on SDS-PAGE gels (e-Pagel, ATTO, Tokyo, Japan). Monoclonal ANTI-FLAG^®^ M2-Peroxidase (HRP) antibody produced in mouse clone M2 (Sigma-Aldrich, Schnelldorf, Germany) and Anti-GFP pAb-HRP-DirecT (MBL, Nagoya, Japan) were used at 1:10,000 dilution to detect FLAG- and GFP-tagged proteins, respectively.

For immunoblot analysis of foxtail millet proteins, the protein concentrations of the SF and EF were determined using Protein Assay Dye Reagent (Bio-Rad) with BSA as a standard. 5 µg proteins were separated on SDS-PAGE gels (e-Pagel, ATTO, Tokyo, Japan). Anti-cFBPase (AS04 043, Agrisera, Vånnås, Sweden) was used as the primary antibody at 1:10,000 dilution. To prepare anti-thaumatin-like protein antiserum, the SGQKPLTLAEFTIGGSQ peptide was synthesized and polyclonal antiserum was raised in rabbit (GenScript, Tokyo, Japan). Anti-rabbit IgG horseradish peroxidase-linked antibody (Promega, USA) was used as the secondary antibody at 1:10,000–20,000 dilution. To detect the signals, Chemi-Lumi One (Nacalai Tesque, Inc., Kyoto, Japan) was used as a substrate for horseradish peroxidase, and the signal was visualized using the ImageQuant LAS 4000 system (GE Healthcare Life Sciences, England).

### RNA and DNA extraction, quantitative PCR

Total RNA was extracted from the samples using a NucleoSpin^®^ RNA Plant kit (Takara Bio Inc., Kusatsu, Japan) in accordance with the manufacturer’s instructions. The RNA was reverse transcribed into cDNA using PrimeScript (Takara Bio Inc.). qPCR was performed using Kapa SYBR FAST qPCR Master Mix (Kapa Biosystems, Roche, Basel, Switzerland) on a QuantStudio 3 Real-time PCR System (Thermo Fisher Scientific, Inc., USA). To quantify *P. palmivora* biomass, total DNA was extracted from the samples as follows. Three days post inoculation with *P. palmivora*, leaves (20 mm in diameter) were ground in liquid nitrogen, dissolved in 300 µl of CTAB (3% cetyltrimethylammonium bromide, 100 mM Tris-HCl, pH 8.0, 1.4 M NaCl, 20 mM EDTA), and incubated at 60 °C for 30 min. The samples were vortexed by adding chloroform, and the water layer was fractionated by centrifugation. The DNA was precipitated with isopropanol, treated with RNase A, and purified using a QIAquick PCR Purification Kit (Qiagen, Venlo, Netherlands) in accordance with the manufacturer’s instructions. qPCR was performed using Kapa SYBR FAST qPCR Master Mix or Luna Universal qPCR Master Mix (NEB) on a QuantStudio 3 system. The primer sequences are listed in Table S7.

### Plasmid construction

All primers used in this study are listed in Table S7. To construct plasmids for qRT-PCR and qPCR analysis, cDNA fragments of *SgJRLs, SgHistone, NbEF1α*, and *PpEF1α* were cloned into pCR™8/GW/TOPO™ or pCR™-Blunt (Thermo Fisher Scientific, Inc.). To analyze virulence and protein accumulation in *N. benthamiana*, cDNA fragments of the signal peptide of *NtPR1a*, the *SgJRL* genes, and *GFP* were amplified and cloned into pCambia-C-3xFLAG (Maqbool et al, 2015).

### Confocal laser-scanning microscopy

Agrobacterium suspensions were infiltrated into the leaves of 4- to 5-week-old *N. benthamiana* plants using a needleless syringe. Three days after infiltration, the leaves were observed under an Olympus Fluoview FV1000 confocal laser-scanning microscope (Olympus, Tokyo, Japan). Plasmolysis was performed by incubating the leaves in 0.75 M mannitol solution. GFP and FM4-64 were excited by a 488-nm laser and detected with bandpass 505–605- and 655–755-nm filters, respectively.

### Preparation and treatment with the INF1 elicitor

The INF1 elicitor was prepared from *Escherichia coli* cells (DH5α) carrying the chimeric plasmid pFB53 containing the *inf1* gene (Kamoun et al. 1997), as previously described (Shibata et al., 2010). *N. benthamiana* leaves were infiltrated with 150 nM INF1 solution using a needleless syringe.

## ACKNOWLEDGEMENTS

We thank S. Kamoun of The Sainsbury Laboratory, Norwich, UK, S. Schornack of The Sainsbury Laboratory, Cambridge, UK, and K. Naito of the National Institute of Agrobiological Sciences for their valuable suggestions for improving the manuscript. This work was supported by JSPS KAKENHI Grant Number 19K06062. We thank the NARO Genebank for providing *Phytophthora palmivora*. Computations were partially performed on the NIG supercomputer at ROIS National Institute of Genetics.

## SUPPORTING INFORMATION LEGENDS

**Fig. S1.**
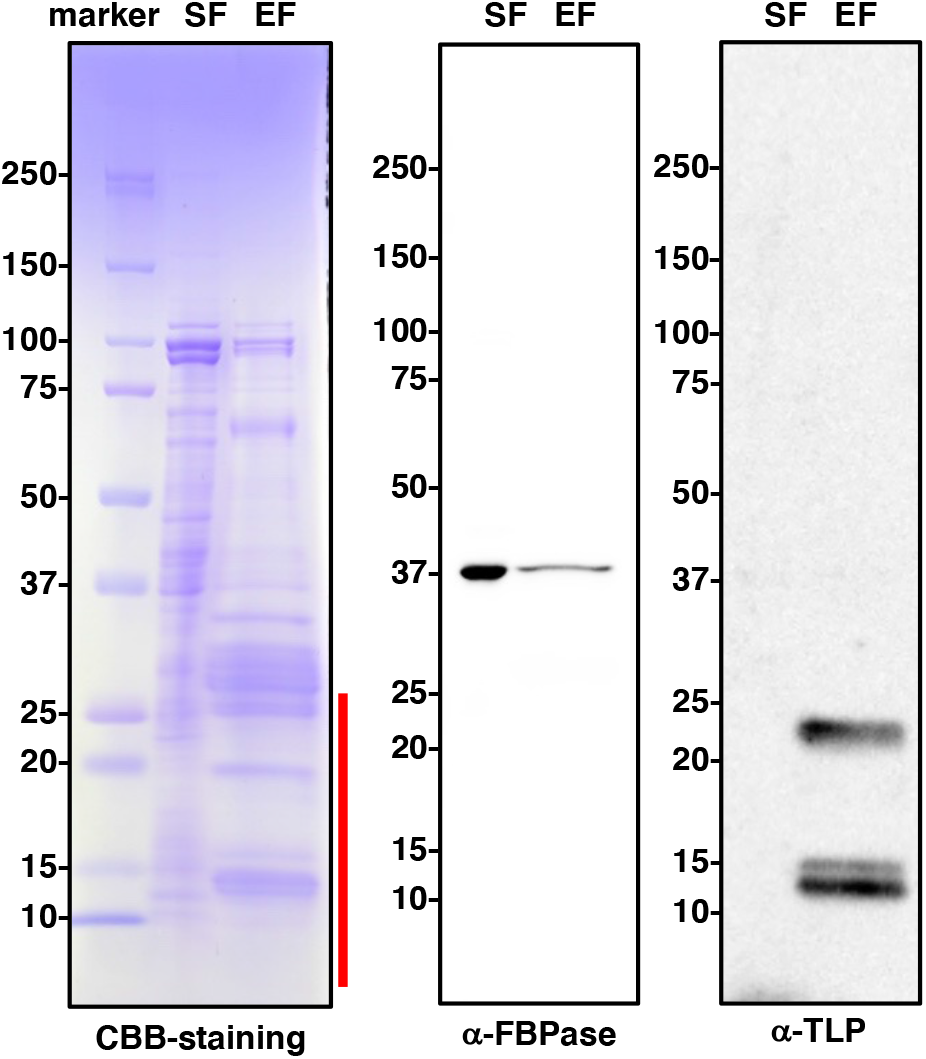
Apoplastic proteins are highly concentrated in the extracellular fluid fraction. Extracellular fluid (EF) and soluble protein fraction (SF) were prepared from foxtail millet leaves 3 days after inoculation with *S. graminicola*. Equal amounts of proteins were separated by SDS-PAGE and analyzed by CBB-staining and immunoblot analysis using anti-FBPase (fructose 1,6-bisphosphatase) and anti-TLP (thaumatin-like protein) antibodies. The red line indicates the range used for the second MS analysis.

**Fig. S2.**
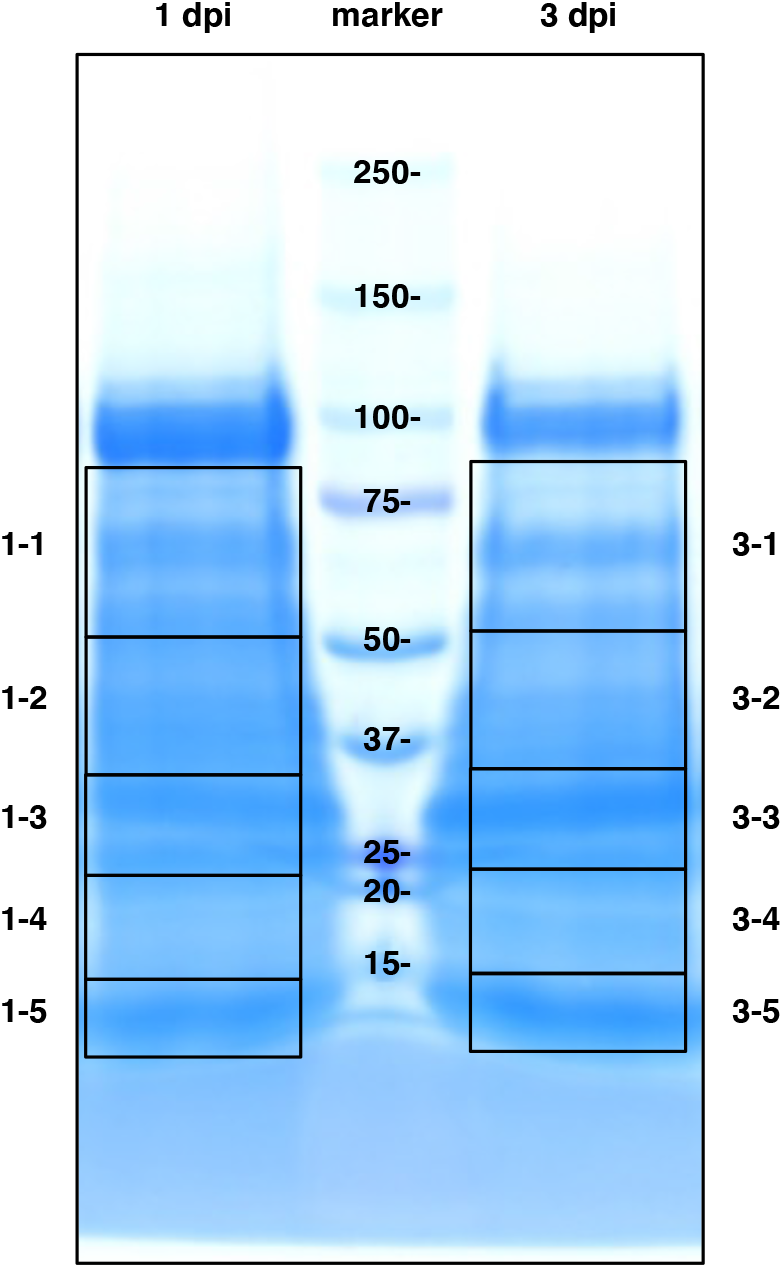
Analysis of extracellular fluid prepared from foxtail millet leaves 1 and 3 days after inoculation with *S. graminicola*. Proteins were separated by SDS-PAGE and analyzed by CBB-staining. Numbered frames indicate samples used for MS analysis.

**Fig. S3.**
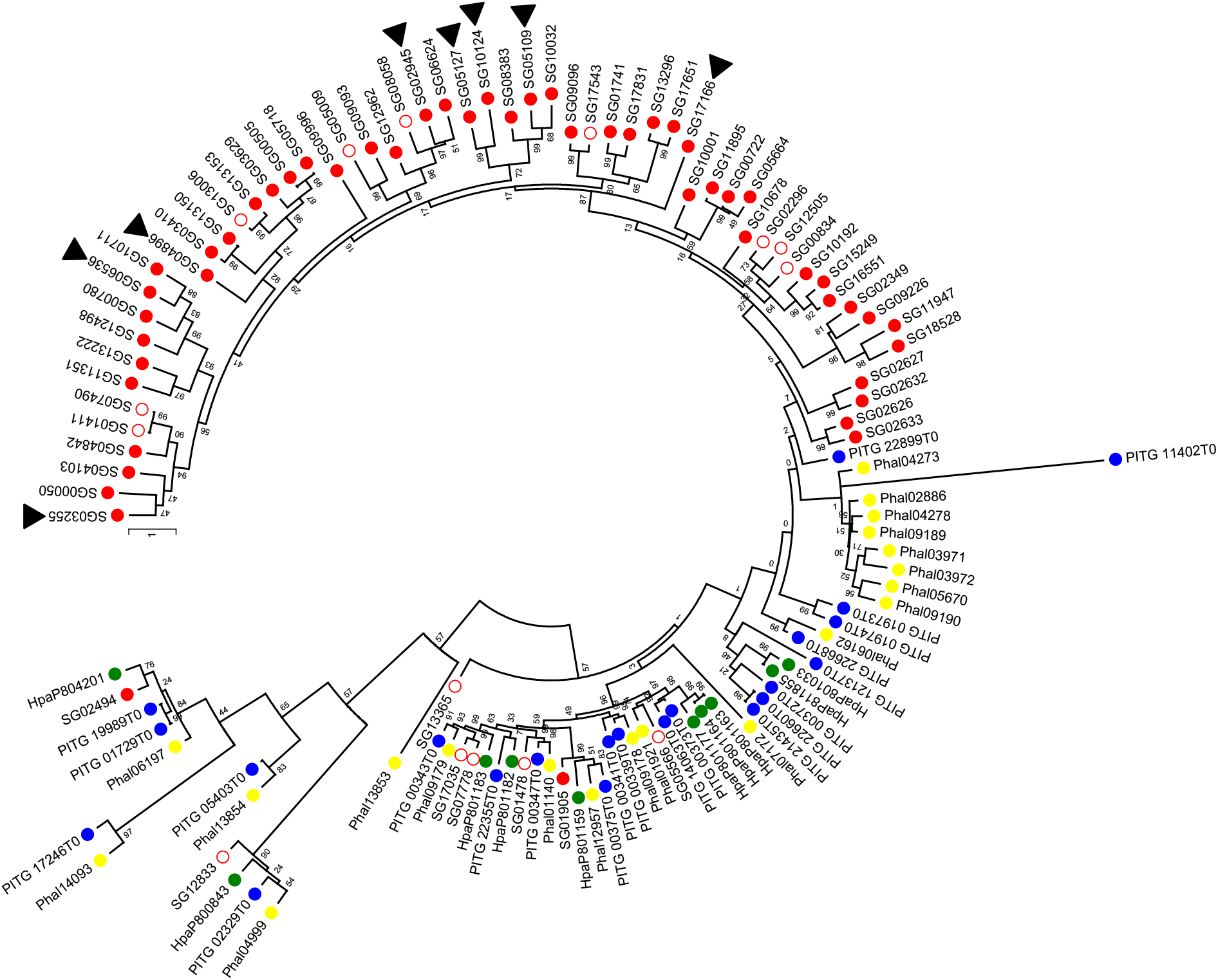
Phylogeny of JRL proteins from *S. graminicola* and the related oomycetes *Hyaloperonospora arabidopsidis, Phytophthora infestans*, and *Plasmopara halstedii*. The tree was constructed using the maximum likelihood method implemented in MEGA6.06-mac, with 1000 bootstrap replicates.

**Fig. S4.**
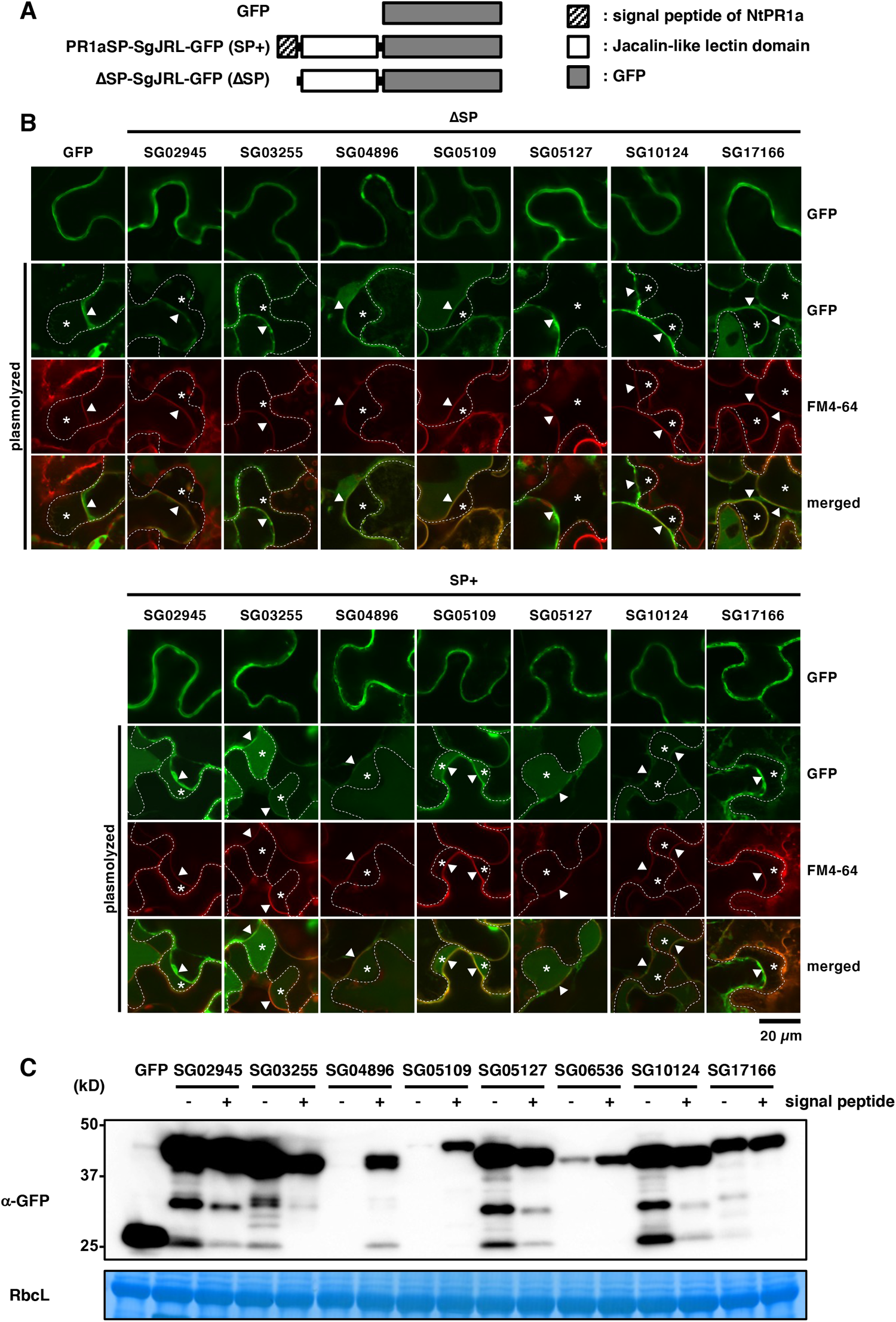
SgJRL fusion proteins with signal peptide accumulate in the plant apoplast. (A) Diagrams of SgJRLs and the fusion proteins used in this study. (B) Genes were transiently expressed in *N. benthamiana* leaves. The leaves were examined 3 days after agroinfiltration. In plasmolyzed cells, the dashed white line marks the cell wall, asterisks indicate the apoplastic space, and triangles indicate the position of the plasma membrane. Bar = 20 µm. (C) Protein accumulation measured by immunoblot analysis using anti-GFP antibody. GFP, PR1aSP-SgJRL-GFP, and ΔSP-SgJRL-GFP were transiently expressed in leaves by agroinfiltration in *N. benthamiana* leaves. Total proteins were extracted from the samples 3 days after agroinfiltration.

**Fig. S5.**
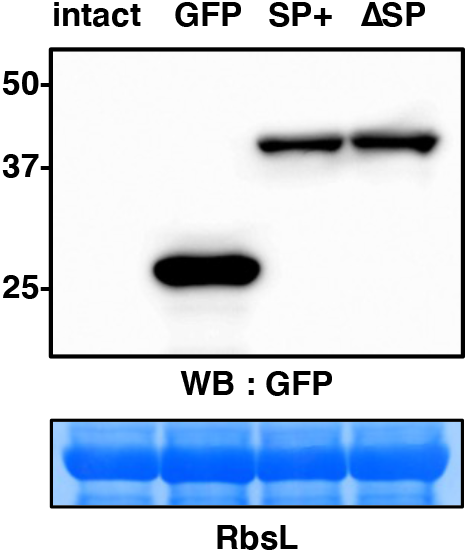
Protein accumulation measured by immunoblot analysis using anti-GFP antibody. GFP, SP-SG06536-GFP, and ΔSP-SG06536-GFP were transiently expressed in leaves by agroinfiltration in *N. benthamiana* leaves. Total proteins were extracted from the samples 2 days after agroinfiltration.

**Fig. S6.**
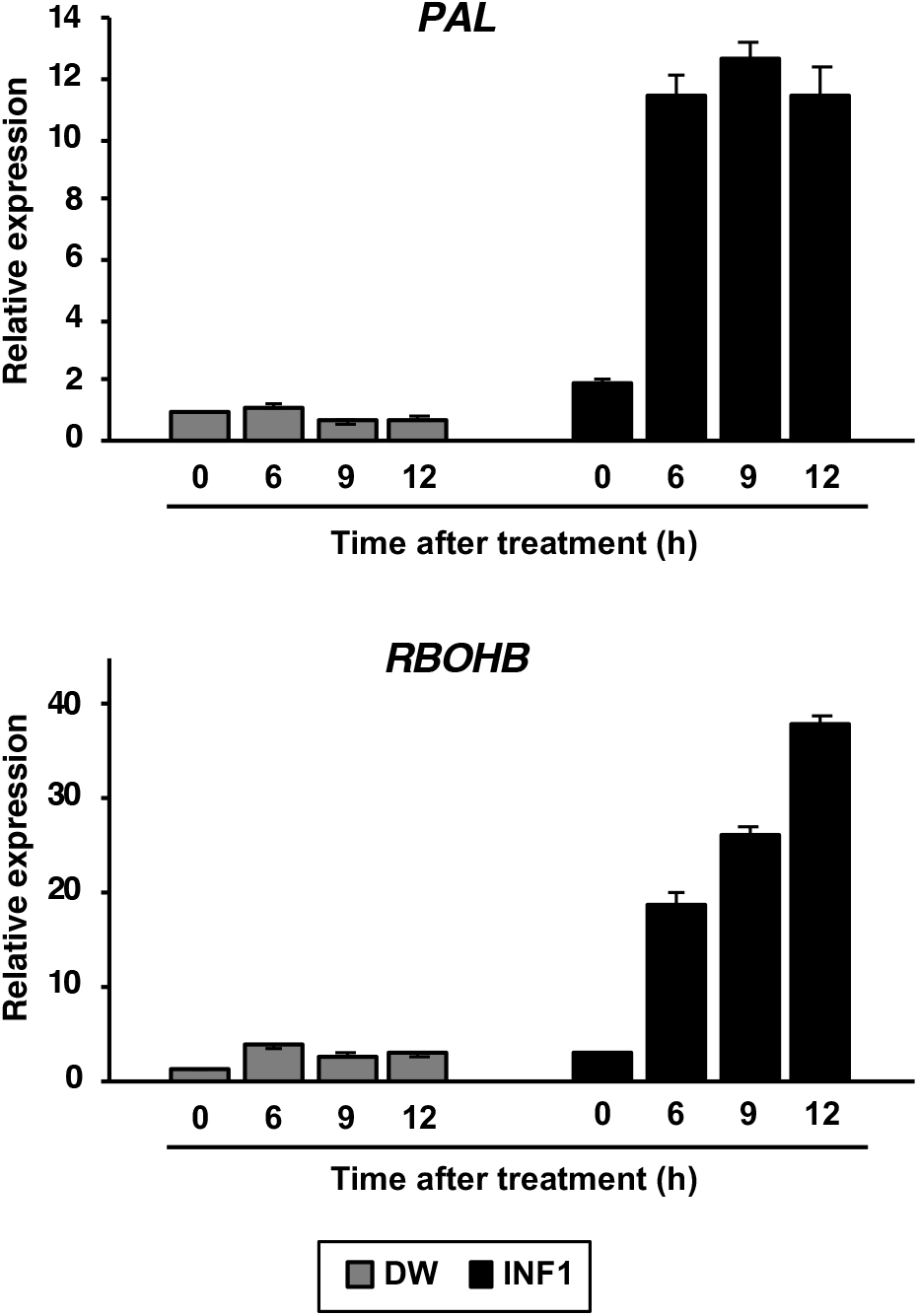
*PAL* and *RBOHB* expression in *N. benthamiana* induced by INF1 peptides. Two days after infiltration of agrobacterium harboring *GFP*, the leaves were subjected to INF1 treatment. Relative expression levels were analyzed by qRT-PCR, with *NbEF1a* used as a reference gene for normalization.

**Table S1** *S. graminicola* proteins in the extracellular fluid from *S. graminicola*-inoculated foxtail millet leaves identified by MS analysis.

**Table S2** *S. graminicola* proteins in the extracellular fluid (EF) and soluble protein fraction (SF) from *S. graminicola*-inoculated foxtail millet leaves identified by MS analysis.

**Table S3** *S. italica* proteins in the extracellular fluid from *S. graminicola*-inoculated foxtail millet leaves identified by MS analysis.

**Table S4** *S. italica* proteins in the soluble fraction (SF) from *S. graminicola*-inoculated foxtail millet leaves identified by MS analysis.

**Table S5** *S. italica* proteins identified in the extracellular fluid (EF) from *S. graminicola*-inoculated foxtail millet leaves by MS analysis.

**Table S6** Jacalin-like domain containing proteins of *Pythium, Magnaporthe*, and *Colletotrichum* characterized by InterProScan analysis.

**Table S7** Primers used in this study.

